# EnGRNT: Inference of gene regulatory networks using ensemble methods and topological feature extraction

**DOI:** 10.1101/2021.08.05.455202

**Authors:** Hakimeh Khojasteh, Mohammad Hossein Olyaee, Alireza Khanteymoori

## Abstract

The development of computational methods to predict gene regulatory networks (GRNs) from gene expression data is a challenging task. Many machine learning methods have been developed, including supervised, unsupervised, and semi-supervised to infer gene regulatory networks. Most of these methods ignore the class imbalance problem which can lead to decreasing the accuracy of predicting regulatory interactions in the network. Therefore, developing an effective method considering imbalanced data is a challenging task. In this paper, we propose EnGRNT approach to infer GRNs with high accuracy that uses ensemble-based methods. The proposed approach, as well as the gene expression data, considers the topological features of GRN. We applied our approach to the simulated Escherichia coli dataset. Experimental results demonstrate that the appropriateness of the inference method relies on the size and type of expression profiles in microarray data. Except for multifactorial experimental conditions, the proposed approach outperforms unsupervised methods. The obtained results recommend the application of EnGRNT on the imbalanced datasets.

## 1. Introduction

Inference of gene regulatory networks (GRNs) is a major challenge in the field of bioinformatics and computational biology. Inference means improving the structure of gene regulatory networks from high throughput expression data. This improvement involves removing noise from the experimental extracted network or adding missed edges to it [1–3]. GRNs are usually represented as graphs in which nodes representing genes and edges represent regulatory interactions between genes. It is possible to determine the level of expression of genes that describe the cellular state in a given condition and time with microarray and other recent technologies such as next generation sequencing. Since genetic or environmental factors can be explained by the risk of a specific disease, finding gene-gene or gene-environment interactions may be critical to gaining a better understanding of the factors affecting the risk of disease [4]. The inference of GRNs to improve their structure has potential implications for medicine and drug design; while finding links between genes through wet-lab experiments is costly and time-consuming [1].

The topology of gene regulatory networks is essential to understand how transcription factors (TFs) regulate genes expression and lead to cellular behaviors such as growth, differentiation, and response to stimuli. Many methods based on machine learning have been developed to infer GRNs, including unsupervised learning methods [5–10], supervised learning methods [11–13], and semi-supervised learning methods [14–18]. Unsupervised methods CLR [5], MRNET [8], and ARACNE [7] are information-theoretic approaches that use mutual information between pairs of genes to reconstruct a gene regulatory network. Other approaches have been proposed based on Mutual Information (MI), including three-way mutual information [19], Conditional Mutual Information (CMI) [20], PCA-CMI [21] method, and finally, the JRAMF [3] method to improve the PCA-CMI method is provided. Unlike unsupervised approaches that exclusively use gene expression data to infer GRNs, supervised learning methods such as SIRENE [13], GENIES [12], and CompareSVM [11], in addition to gene expression data, require known regulatory interactions between TF and target gene to train a model. Semi-supervised methods are the interstellar state of these two machine learning approaches and with special methods that include training a model with labeled and unlabeled data. A comprehensive analysis of these three machine learning methods for inferring GRNs by Maetschke et al. [22] concluded, that in most cases, supervised learning methods have better performance than other methods.

In this paper, we propose a supervised learning-based method called EnGRNT to infer GRNs that is different from previous works in two respects. First, for each TF, we consider the GRN inference as a binary classification problem. In our problem, the number of regulatory relationships known between TF and target genes (class +1) is much less than the number of regulatory relationships specified between TF and the target gene does not exist (class −1) [23]. Hence, we deal with imbalanced data classification. Some previous approaches have ignored this challenge and have obtained high accuracy by classifying the majority of samples as negatives [11]. In this work, we use the underbagging [24] ensemble learning algorithm to address the imbalance problem. Using this method, we created various bootstraps for each TF and a classifier is trained with each bootstrap. All local models ultimately create an ensemble inference engine that integrates them which can predict new interactions in the network and even if necessary, modify the gene regulatory network. Second, Fire et al. [25] shown that topological features can be used to increase the accuracy of link prediction in social network analysis. In this work, we extend their techniques to solve an important bioinformatics problem, i.e. GRN inference. For this aim, a feature extraction phase is performed, which extracted the set of topological features of regulatory networks. Therefore, EnGRNT is a supervised learning method that uses the gene expression data and topological features extracted from GRN to train ensemble classifiers and ultimately improves the GRN structure. To investigate the proposed approach against the known real network, the evaluation was performed on simulated data and stable state of Escherichia coli (E. coli) [23]. Furthermore, three state-of-the-art information-theoretic network inference methods, namely including CLR [5], MRNET [8] and ARACNE [7] are applied to the simulated E. coli dataset. According to the obtained results, our approach outperforms previous unsupervised approaches, and the application of topological features combined with gene expression data in model training has improved the performance of the inference method.

The rest of the paper is organized as follows. Section 2 firstly provides a brief review of the proposed method. In the following, the proposed method is described in detail. Section 3 introduces three unsupervised learning methods that in this study have been also employed to infer GRN. Data Set, performance metrics, and results are presented in section 4. Finally, Section 5 summarizes the conclusions.

## 2. Method

### An overview of proposed method

The outline of the proposed approach is shown in Figure 1, which consists of four basic phases:

1. Extracting biological knowledge: The GRN inference is considered as a binary classification problem and it needs to extract a series of biological knowledge to train classifiers. First, we should extract all TFs in GRN and their corresponding target genes in different biological experimental conditions.
2. Topological feature extraction: In this phase, a set of topological features is extracted specific to regulatory networks. GRNs have certain topological properties that affect the process of regulating genes. These networks have high degree centrality nodes that act as regulators for many of genes. Also, these networks have locality properties that reflect specific performance in the network.
3. Training ensemble classifiers for each TF: The biological knowledge extracted from phase 1 as well as the obtained topological features in phase 2, are used to train ensemble classifiers. Since these training sets are highly imbalanced, it is necessary to tackle this challenge. In this regard for each TF, various bootstraps are created by UnderBagging algorithm. Next, for each TF, an ensemble of binary classifiers is trained based on the gene expression data and extracted topological features.
4. Predicting new interactions and reconstructing gene regulatory network: At the last phase, new interactions in the regulatory network are inferred. Finally, the outputs obtained by ensemble of classifiers determines that the existing network needs to be modified or some interactions should be removed.

**Fig.1.**
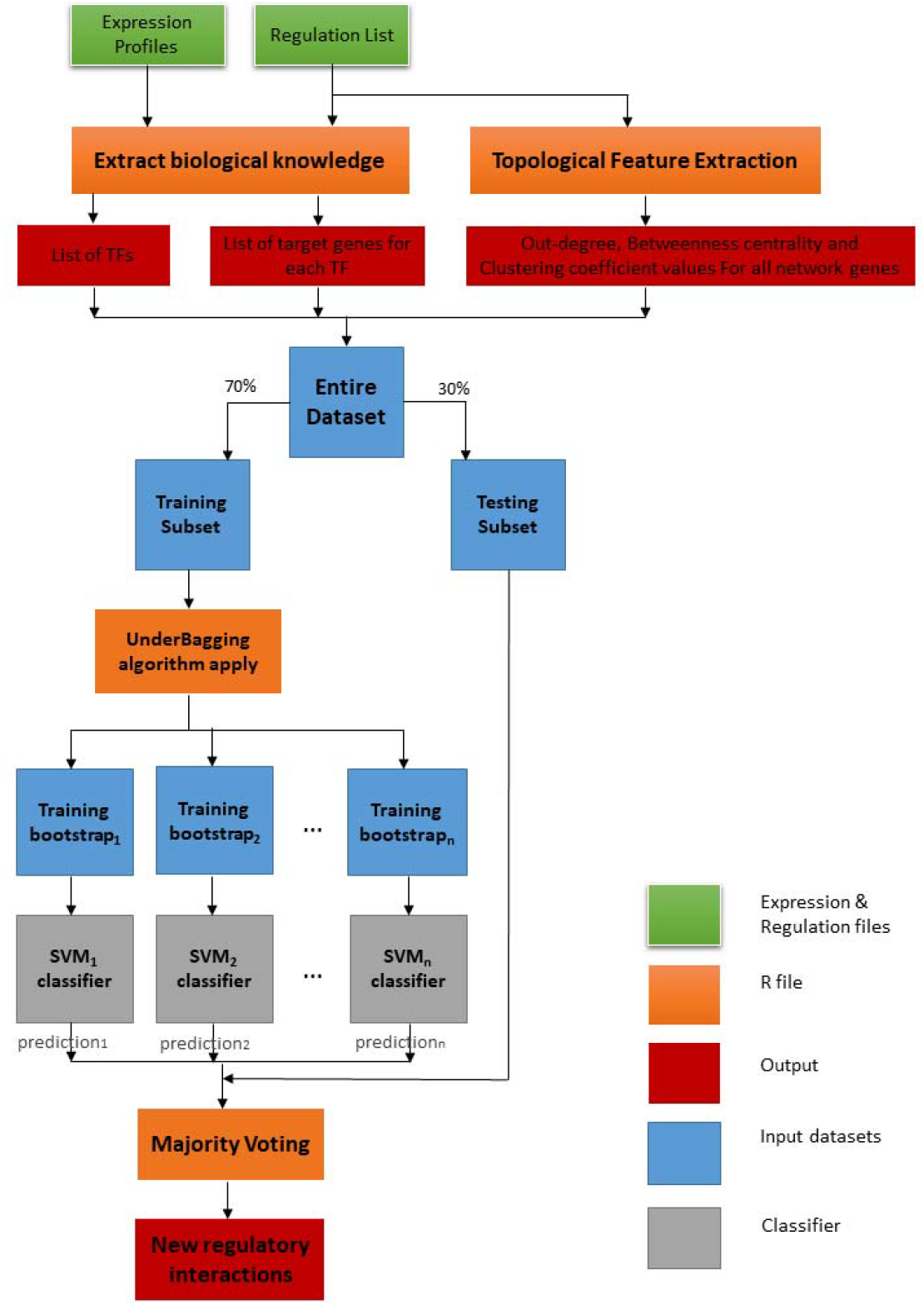
The workflow of proposed method

#### 2.1. Extracting biological knowledge

To reconstruct GRN, we need first to extract all TFs in GRN and their corresponding target genes. For this purpose, two types of inputs are required: first, the list of genes and their expression values for experimental conditions (knockdown, knockout and multifactorial), which is a vector of expression values in a compendium of expression profiles for a given experimental condition. Second, the list of known regulation relationship between TFs and target genes. Such lists can usually be established from available databases of experimentally regulations, e.g. RegulonDB [23] for E. coli genes.

#### 2.2. Topological feature extraction

The gene regulatory networks are considered as directed graphs. If we consider only the genes coded for TFs, we can simply imagine a network whose nodes, generators of TFs and links in that network correspond to the regulatory interactions between the genes. Such a network (or a graph) naturally has directional edges, for example, activating the Y gene by a genetic regulation product X is a completely different process of activation of X by the product Y [26].

In the context of gene regulatory networks, several topological properties are considered: (1) Each gene in GRN is usually influenced by a small number of other genes; the distribution resulted by incoming interactions is narrow; (2) A few genes (called hubs) regulate a large number of other genes; hence the distribution resulted by outgoing interactions is widespread, possibly a power law; (3) The regulatory networks are robust to internal fluctuations (stochastic nature of protein production) or external signals (variations in temperature, oxygen levels, nutrient abundance). This property is also found at many levels of other biological organizations. (4) Regulatory networks show modular structure and in particular, certain subgraphs (network motifs) are presented in comparison with randomized networks [26].

Based on the topological properties of GRN, the following features can be extracted:

##### Out-degree 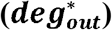

The topology of gene regulatory networks is dominated by hub nodes, which have higher out-degree than other nodes in the network. Based on the knowledge of these networks, it can be found that these nodes are the same transcription factors (TFs) that regulate target genes. Suppose a GRN including n nodes (genes), for all nodes the normalized out-degree is calculated according to equation (1):

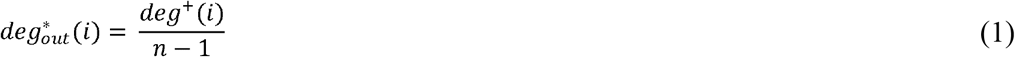

##### Betweenness centrality (*C_B_*)

The hub nodes in GRN act as network connectors, which means connecting a part of the network with another part. This feature is called betweenness centrality [27]. This feature is often used to find nodes that act as bridges from part of the graph to another. The betweenness centrality is calculated according to the following equation for all network nodes:

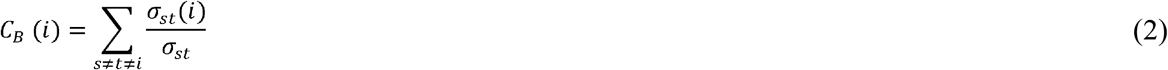

Where *σ_st_* is the total number of shortest paths from node s to node t and *σ_st_* (*i*) is the number of shortest paths that pass from node *i*. The betweenness centrality increases with the number of vertices in the network, so its normalized version is often considered to be centered between zero and one. The normalized betweenness centrality is calculated by dividing equation (2) to number of possible edges in the network regardless of the node *i*. Therefore, the normalized betweenness centrality for all network nodes is calculated as follows:

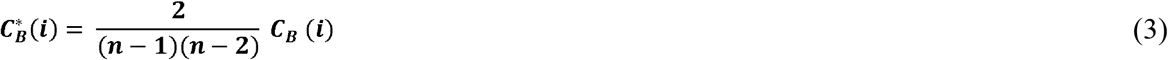

##### Clustering coefficient (*C_i_*)

In graph theory, the clustering coefficient is a measure of the degree to which nodes in a graph tend to form a cluster together. This common property is called modularity, which means that the entirety of the gene network consists of subnets of recognizable genes or modules, each of which corresponds to the function in the network. Therefore, the clustering coefficient is another feature that can be extracted from these networks. In addition, Wu et al. [28] proposed a degree-related clustering coefficient for link prediction in complex networks. Unlike the classic clustering coefficient, the new coefficient is very robust, especially outperformed on sparse networks with low average clustering coefficients. Therefore, the clustering coefficient for all network nodes is calculated from the following equation:

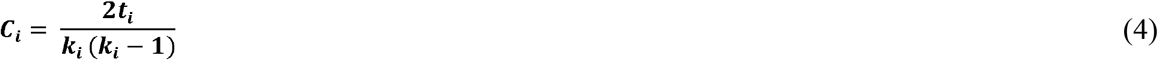

Where *t_i_* is the number of triangles passing through node *i* and *k_i_* is the degree of node *i*.

#### 2.3. Training ensemble classifiers for each TF

In this section, we briefly introduce the class imbalance problem. In classification, a dataset is said to be imbalanced when the number of instances which represents one class is smaller than the ones from other classes. The imbalance problem occurs when a class (minor or positive class) has a small sample number in the dataset. Concerning the inference problem, the number of regulatory relationships known between TF and target genes is much less than the number of regulatory relationships specified between TF and the target gene does not exist. On account of the importance of the imbalance problem, a category of techniques has been developed to overcome this challenge [29]. In the proposed approach, we focus on ensemble-based methods which consist in the manipulation of the training examples in such a way that each classifier is trained with a different training set.

Ensemble learning methods such as bagging [24] and boosting [30] have traditionally provided promising results to improving the prediction accuracy in classification. These techniques have been widely used in several areas and applications to progress in computational efficiency. Bagging [24] which stands for bootstrap aggregation obtains diversity in classification by randomly sampling the data – with replacement – from the entire training dataset. Each sample is used to train classifiers that can be of the same type or different types. Simple majority vote is used to fuse the resultant models. Boosting [30] is similar to the ensemble learning method of bagging in that it resamples the data, however, it strategically samples the subset of the training data to create several weak classifiers. The classifiers are combined through an n-way majority voting. For example, if there are three classifiers, the first subset of the training data is selected randomly, the second subset is selected in an informative way as per the boosting algorithm. The third subset is sampled based on the instances where both the first and second classifier disagrees with each other. Hence, during fusion of data, a strong classifier is created.

Many methods have been proposed using bagging ensembles to deal with class imbalance problems due to its simplicity and good generalization ability. The hybridization of bagging and data preprocessing techniques is usually simpler than their integration in boosting. In these methods, the key factor is the way to collect each bootstrap replica, that is, how the class imbalance problem is considered to obtain a useful classifier in each iteration without ignoring the importance of the diversity [29]. One of the main algorithms in the bagging family is the UnderBagging algorithm, which we use to address the class imbalance problem in GRN.

In phase 3 of the proposed approach, the combination of phase 1 and phase 2 yields constitute the training set. First, the training set is divided into two training and test subsets for each TF. By applying the UnderBagging algorithm, various bootstraps are created and a classifier is trained with each bootstrap. For each TF, an ensemble of binary classifiers is trained based on gene expression data and topological features extracted by two data classes: the genes that are known to be regulated by that TF and the genes that are not known to be regulated by that TF. Since we do not have enough knowledge about relationship between the level of expression of TF measurement and its target genes, we suppose that two genes likely exhibit similar expression patterns, if they are regulated by the same TF [11, 13].

For each local model associated with each TF, Support Vector Machine (SVM) is used as an ensemble classifier. The reason for using this classifier is that it has been successfully applied to GRN inference [11, 13, 22]. In the proposed approach, as a kernel for SVM, Gaussian kernel is used. Because it was successfully used in SIRENE [13] as well as CompareSVM [11] and they discussed that Gaussian kernel generally has the best performance for all experimental conditions compared to other kernels. In addition, for data such as microarray where the number of samples is much less than the number of features, choosing Gaussian kernel seems more logical [31]. It should be noted that for more investigation, the sigmoid kernel is also implemented for SVM to compare their performance.

#### 2.4. Reconstruction of gene regulatory network

In phase 4 of the proposed approach, new interactions from the regulatory network are identified. By using the local models created for each TF, the list of new genes can be assigned to the local model of each TF. If scores obtained from classifiers meet threshold value, then a majority vote will be taken between predictions of these classifiers and finally it will be determined whether the new gene is regulated by that TF. The final phase is to combine all scores of the local models to rank the candidate TF-gene interactions by descending scores.

## 3. Unsupervised methods

This section reviews some unsupervised learning methods for network inference which are based on information-theoretic concepts. A number of unsupervised learning methods have been utilized to infer GRNs. Unlike supervised approaches, unsupervised inference algorithms apply to only gene expression profiles for gene network construction. For comparison, we also performed three state-of-the-art information-theoretic network inference methods on the compendium data. Information-theoretic approaches identify candidate interactions by estimating pairwise gene expression profile mutual information (*M_ij_*) [32]. This criterion measures the degree of dependence between the two genes *X_i_* and *X_j_*:

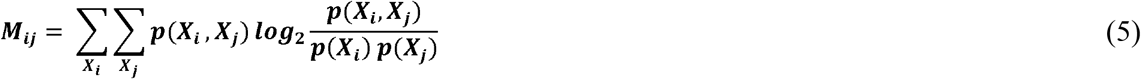

where *p*(*X_i_*, *X_j_*) the probability distribution functions of *X_i_* and *X_j_*, and *p*(*X_i_*) and *p*(*X_j_*) are the marginal probability distribution functions of *X_i_* and *X_j_*,respectively.

The MRNET (The Minimum Redundancy NETworks) method reconstructs the network using a feature selection technique called redundancy reduction-enhancement (Minimum Redundancy Maximum Relevance), which is based on measuring the mutual information (Mij) between genes[8]. To infer a network, a feature selection algorithm for each gene (as the target gene) is performed on a set of the rest of the genes. These genes are considered to be regulators of that target gene.

Relevance network (RELNET) is the simplest method based on mutual information [33]. For each gene pair, mutual information (*M_ij_*) is estimated, and the interaction is created between genes *i* and *J* if the mutual information is greater than a threshold. The CLR algorithm is an extension of RELNET [5]. This method computes a score related to the experimental distribution of mutual information values. In this method, for each gene *i* and all other genes *J* ≠ *i*, a maximum z-score is calculated as follows:

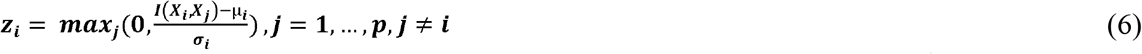

Where *μ_i_* and *σ_i_* are the mean and standard deviation of gene *i* and *I*(*X_i_*, *X_j_*) the mutual information between the two genes. Finally, the weight of the interaction between the two genes *i* and *J* is obtained as follows:

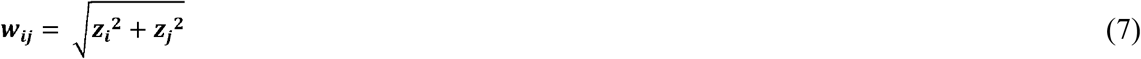

The ARACNE algorithm (Algorithm for The Reconstruction of Accurate Cellular Networks) is another extension of RELNET, which applies Data Processing Inequality (DPI) to filter indirect interactions [7]. Based on DPI, if gene *i* interacts with gene *J* through gene *k*, there is the following inequality:

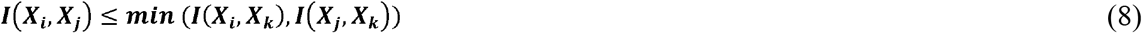

This algorithm considers each triple of edges independently, and calculates the mutual information (*MI*) for each gene pair within the triplet. So, the weakest edge of each triplet is interpreted as an indirect interaction and is removed.

## 4. Experimental results

### 4.1. Data Set

There are usually two methods for generating GRN benchmark data: (1) employing the networks that obtained from cultivate inside the living creature. (2) using simulated artificial networks in which the expression data is used to make perturbation artificially by some kind of computer simulation [34]. The proposed approach was tested on simulated E. coli data extracted by GeneNetWeaver [35] from different nodes (number of genes) in the sizes (10, 30, 50,100,150,200 and 500) of the genes and types of experimental conditions (knockdown, knockout, and multifactorial).

GeneNetWeaver [35] is an open source tool for benchmarking silico and profiling for network inference, which applies the real microarray compendia (derived from the first method) along with synthetic data. This tool simulates and generates datasets for gene expression profiles in a variety of knockdown, knockout, and multifactorial for E. coli. In knockdown experiment, expression of one or more genes are reduced. In knockout experiment, one of the gene expressions is entirely made inactive and in multifactorial experiment, a few number of genes expression values are perturbed though a tiny random number.

### 4.2. Performance Metrics

In order to evaluate the performance of different inference methods to predict the regulatory interactions between genes, the GRN algorithm generates a ranked list of predicted edges. This list is sorted according to the confidence in the predictions, so that the first edge in the list corresponds to the edge with the highest confidence. By considering both directions, the number of possible edges in a network with N-genes is N(N - 1). From a ranked edge-list of this kind, an interconnected network of k edges is obtained by setting a cutoff k which considers the first k edge as the present and the rest of the edges are considered as absent. Therefore, k is a parameter that controls the number of edges in the predicted network [26].

The evaluation metric is a key factor in evaluating the classification performance. One of the metric that demonstrates the efficiency of GRN inference method is using of the Receiver Operating Characteristic (ROC) curve [36]. To draw a ROC curve, only the true positive rate (TPR) and false positive rate (FPR) are needed. The TPR defines how many true positive results occur among all positive samples available during the test. On the other hand, FPR, defines how many false positive results occur among all negative samples available during the test. A ROC space is defined by FPR and TPR as x and y axes, respectively, which depicts relative trade-offs between true positive (benefits) and false positive (costs). To evaluate the quality of the proposed approach and compare it with other methods, AUC (area under the Receiver Operator Characteristics curve) has been used. AUC is a metric that demonstrates the quality of inference method. The desired AUC for classification is bounded to 0.5 and 1 (AUC E (0.5,1)). When AUC is close to 0.5, indicating that prediction is no better than accidental guessing, and AUC close to 1 represents a very accurate prediction [37].

### 4.3. Results

To evaluate the proposed approach, we have investigated 21 sub networks related to E. coli. In order to demonstrate the performance, proposed approach without topological feature extraction phase (called EnGRN) is also performed. For better evaluation, the proposed approach has been compared with three common unsupervised methods including ARACNE, CLR and MRNET in the field of gene regulatory networks construction. We used SVM library LIBSVM developed by Chang et al. [38] for implementation of proposed approach. Unsupervised methods are performed by the minet package [37] in R and with Spearman’s adjacency matrix and default parameters.

We evaluated prediction accuracy of EnGRNT using two different kernels in microarray simulated data. We have tested EnGRNT on simulated E. coli data extracted by GeneNetWeaver [35] of different nodes (number of genes) ranging from 10 to 500 in three different biological experimental conditions. Table 1 to 4 show the AUC values for different methods in three experimental conditions (knockdown, knockout, and multifactorial) and also average of three experimental conditions (all).

**Table 1.** Prediction accuracy (AUC) of unsupervised and supervised methods on knockdown experimental condition.

| Size | ARACN<br>E | CLR<br>(Spearman<br>) | MRNET | EnGRN<br>(Sigmoid) | EnGRN<br>(Radial) | EnGRNT<br>(Sigmoid) | EnGRNT<br>(Radial) |
| --- | --- | --- | --- | --- | --- | --- | --- |
| 10 | 0.2845 | 0.3054 | 0.2796 | 0.75 | 0.625 | 0.75 | **0.75** |
| 30 | 0.3031 | 0.4019 | 0.3984 | 0.683 | 0.6071 | 0.6806 | **0.6458** |
| 50 | 0.3123 | 0.4395 | 0.457 | 0.6635 | 0.6093 | 0.6319 | **0.6584** |
| 100 | 0.368 | 0.5001 | 0.4985 | 0.5577 | 0.5498 | 0.5744 | **0.6078** |
| 150 | 0.3389 | 0.4773 | 0.482 | 0.545 | 0.5297 | 0.5603 | **0.6047** |
| 200 | 0.2878 | 0.4484 | 0.4273 | 0.5433 | 0.5274 | 0.5482 | **0.6302** |
| 500 | 0.3028 | 0.4722 | 0.4689 | 0.5083 | 0.5022 | 0.5391 | **0.5682** |

**Table 2.** Prediction accuracy (AUC) of unsupervised and supervised methods on knockout experimental condition.

| Size | ARACNE | CLR<br>(Spearman<br>) | MRNET | EnGRN<br>(Sigmoid) | EnGRN<br>(Radial) | EnGRNT<br>(Sigmoid) | EnGRNT<br>(Radial) |
| --- | --- | --- | --- | --- | --- | --- | --- |
| **10** | 0.4742 | 0.4995 | 0.4764 | 0.75 | 0.75 | 0.9667 | **0.6667** |
| **30** | 0.2986 | 0.3248 | 0.3332 | 0.6667 | 0.6375 | 0.6562 | **0.6667** |
| **50** | 0.3332 | 0.4831 | 0.4808 | 0.6634 | 0.703 | 0.6467 | **0.6516** |
| **100** | 0.376 | 0.4976 | 0.4834 | 0.5589 | 0.572 | 0.5702 | **0.6018** |
| **150** | 0.3463 | 0.4678 | 0.4665 | 0.5451 | 0.5714 | 0.5607 | **0.6052** |
| **200** | 0.305 | 0.4461 | 0.4422 | 0.5319 | 0.5547 | 0.5379 | **0.6504** |
| **500** | 0.2934 | 0.4606 | 0.4559 | 0.5143 | 0.5112 | 0.5345 | **0.596** |

**Table 3.** Prediction accuracy (AUC) of unsupervised and supervised methods on multifactorial experimental condition.

| Size | ARACNE | CLR<br>(Spearman<br>) | MRNET | EnGRN<br>(Sigmoid) | EnGRN<br>(Radial) | EnGRNT<br>(Sigmoid) | EnGRNT<br>(Radial) |
| --- | --- | --- | --- | --- | --- | --- | --- |
| **10** | 0.5636 | 0.5938 | 0.574 | 0.75 | 0.875 | 0.7919 | **0.6667** |
| **30** | 0.327 | 0.4307 | 0.4342 | 0.6375 | 0.625 | 0.6389 | **0.6429** |
| **50** | 0.3732 | 0.6686 | 0.6667 | 0.6131 | 0.7119 | 0.5901 | **0.6672** |
| **100** | 0.4572 | 0.5856 | 0.5649 | 0.5602 | 0.5477 | 0.568 | **0.6636** |
| **150** | 0.4007 | 0.7902 | 0.7555 | 0.5317 | 0.5276 | 0.5504 | **0.6134** |
| **200** | 0.4027 | 0.7162 | 0.7053 | 0.5265 | 0.5305 | 0.547 | **0.628** |

**Table 4.** Prediction accuracy (AUC) of unsupervised and supervised methods on all (average of three experimental conditions).

| Size | ARACNE | CLR<br>(Spearman<br>) | MRNET | EnGRN<br>(Sigmoid) | EnGRN<br>(Radial) | EnGRNT<br>(Sigmoid) | EnGRNT<br>(Radial) |
| --- | --- | --- | --- | --- | --- | --- | --- |
| **10** | 0.4407 | 0.4662 | 0.4433 | 0.75 | 0.75 | 0.8194 | **0.6944** |
| **30** | 0.3095 | 0.3858 | 0.3886 | 0.6624 | 0.6232 | 0.6585 | **0.6518** |
| **50** | 0.3395 | 0.5304 | 0.5348 | 0.6466 | 0.6747 | 0.6229 | **0.659** |
| **100** | 0.4004 | 0.5278 | 0.5150 | 0.5589 | 0.5565 | 0.5708 | **0.6244** |
| **150** | 0.3619 | 0.5784 | 0.568 | 0.5406 | 0.5429 | 0.5571 | **0.6077** |
| **200** | 0.3318 | 0.5369 | 0.5249 | 0.5339 | 0.5375 | 0.5443 | **0.6362** |

In order to better illustrate the improvement of the proposed approach, in Figures 2–5, the AUC values are plotted for each 21 sub networks in three experimental conditions (knockdown, knockout, and multifactorial) and also average of three experimental conditions (all). As can be seen in Figure 2, the proposed approach using both kernels (Gaussian and sigmoid) on average has higher AUCs than the unsupervised methods. According to Figures 2–5, with the exception of multifactorial experimental condition, prediction accuracies of unsupervised methods (CLR, MRNET and ARACNE) are low in general.

**Fig. 2.**
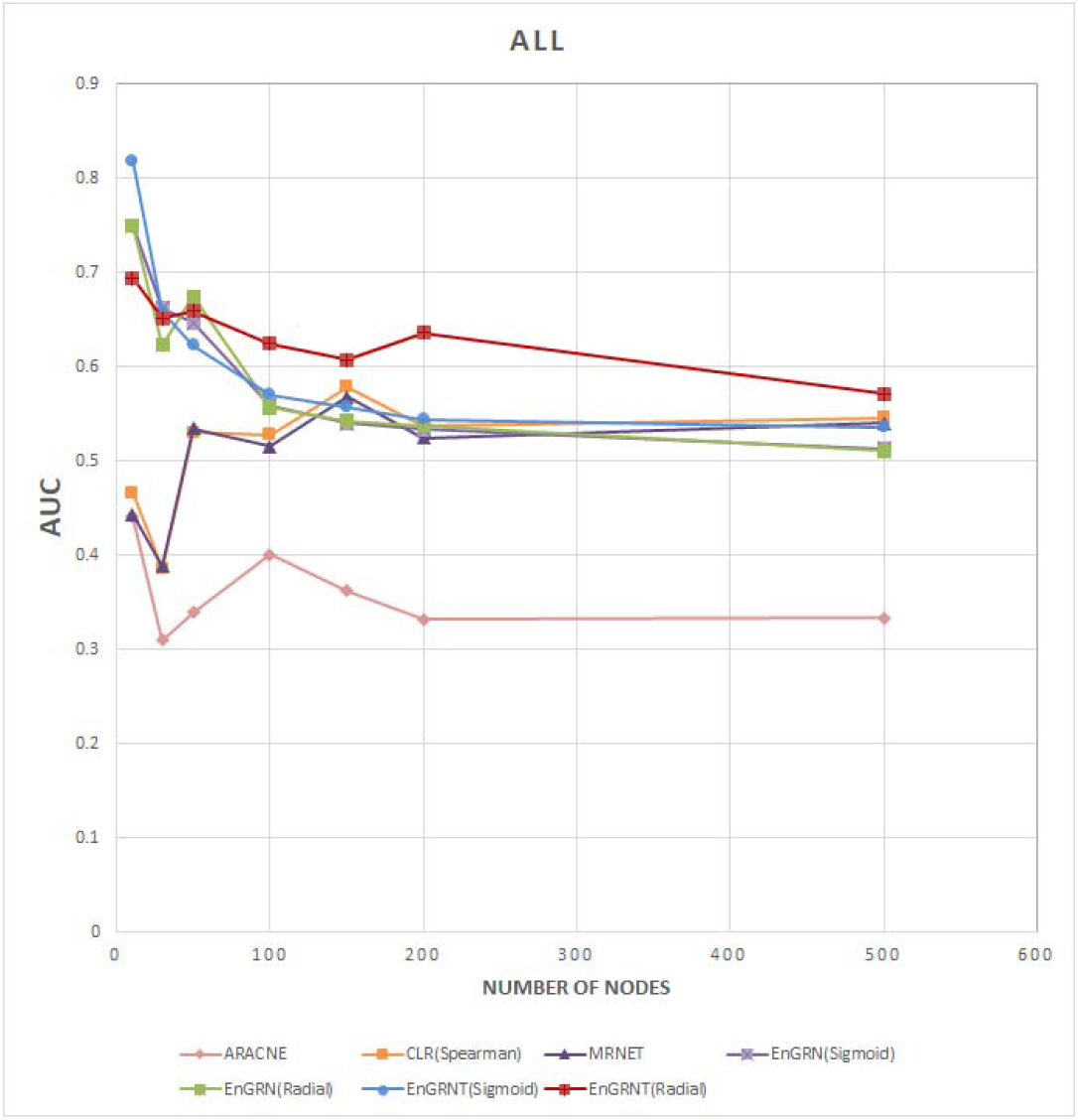
Prediction accuracy (AUC) of unsupervised and supervised methods on all (average of three experimental conditions)

In the following, we first evaluate the prediction accuracy of unsupervised methods. In unsupervised methods, CLR (Spearman) and MRNET had similar results for all biological conditions. The ARACNE performed very poorly as shown in Figures 2–5. For knockdown and knockout experimental condition, CLR (Spearman) and MRNET had poor performance in all experimental conditions and their performance were not better than random guess, but in multifactorial experimental condition, they were able to perform better than supervised methods on the networks with (≥150) nodes. As a result, with the exception of multifactorial experimental condition, unsupervised methods had very low accuracy in GRN inference.

Our investigations demonstrate good prediction accuracies for proposed approach on all experimental conditions. As can be seen in Figure 3 and Figure 4, the proposed approach has higher prediction accuracies than other methods in knockdown and knockout experimental condition. In comparison with unsupervised methods, the accuracy of the proposed approach using the sigmoid kernel exceeds even 90% in knockout experimental condition for a network with 10 nodes. In multifactorial experimental condition, the proposed approach represented higher prediction accuracies than EnGRN in both kernels for networks with (≤ 100) nodes, but as expected, prediction performance using Gaussian kernel was better than sigmoid kernel in most cases as shown in Figure 5 CLR (Spearman) and MRNET methods outperform supervised methods in larger networks.

**Fig.3.**
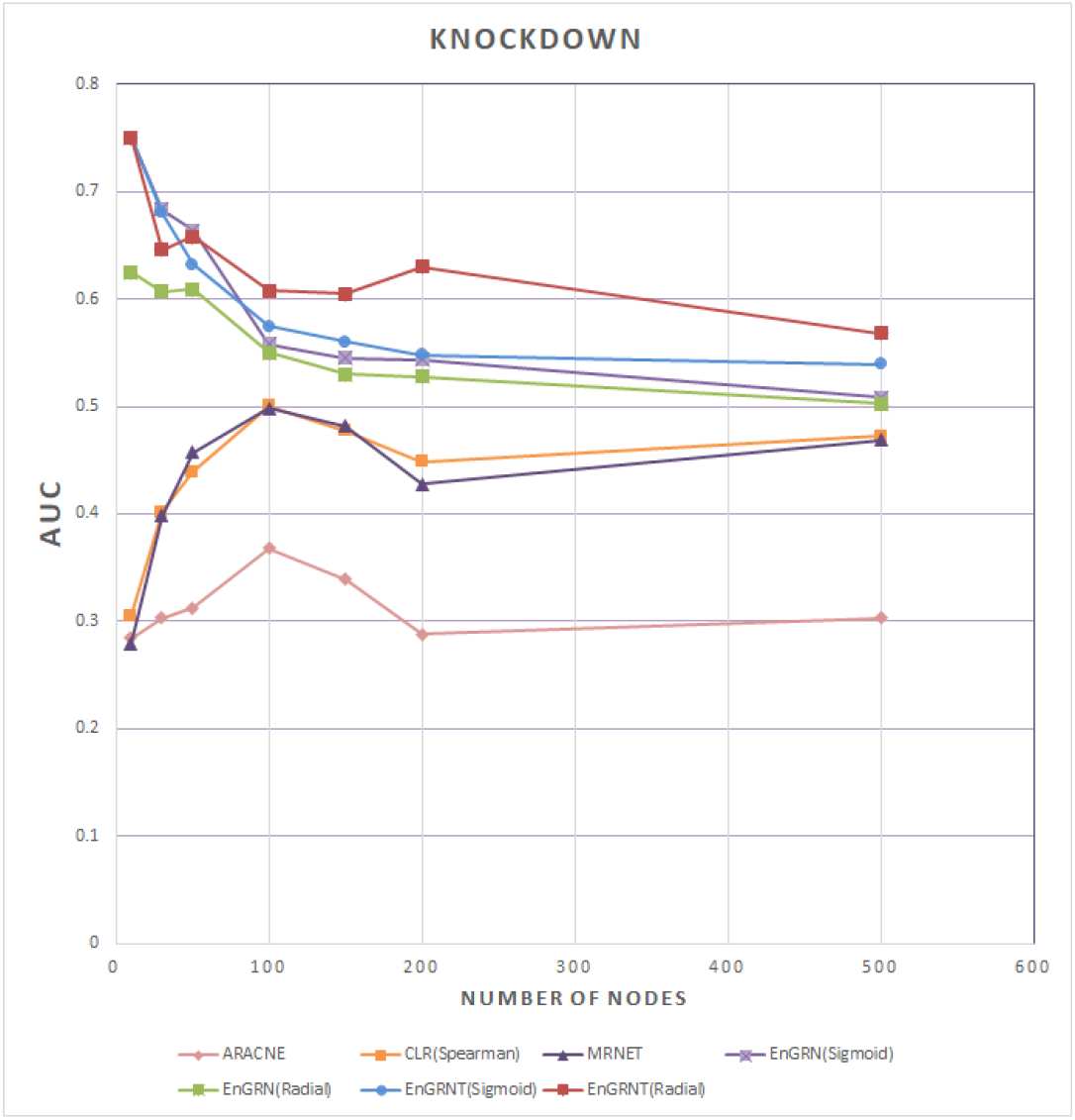
Prediction accuracy (AUC) of unsupervised and supervised methods on knockdown experimental condition

**Fig.4.**
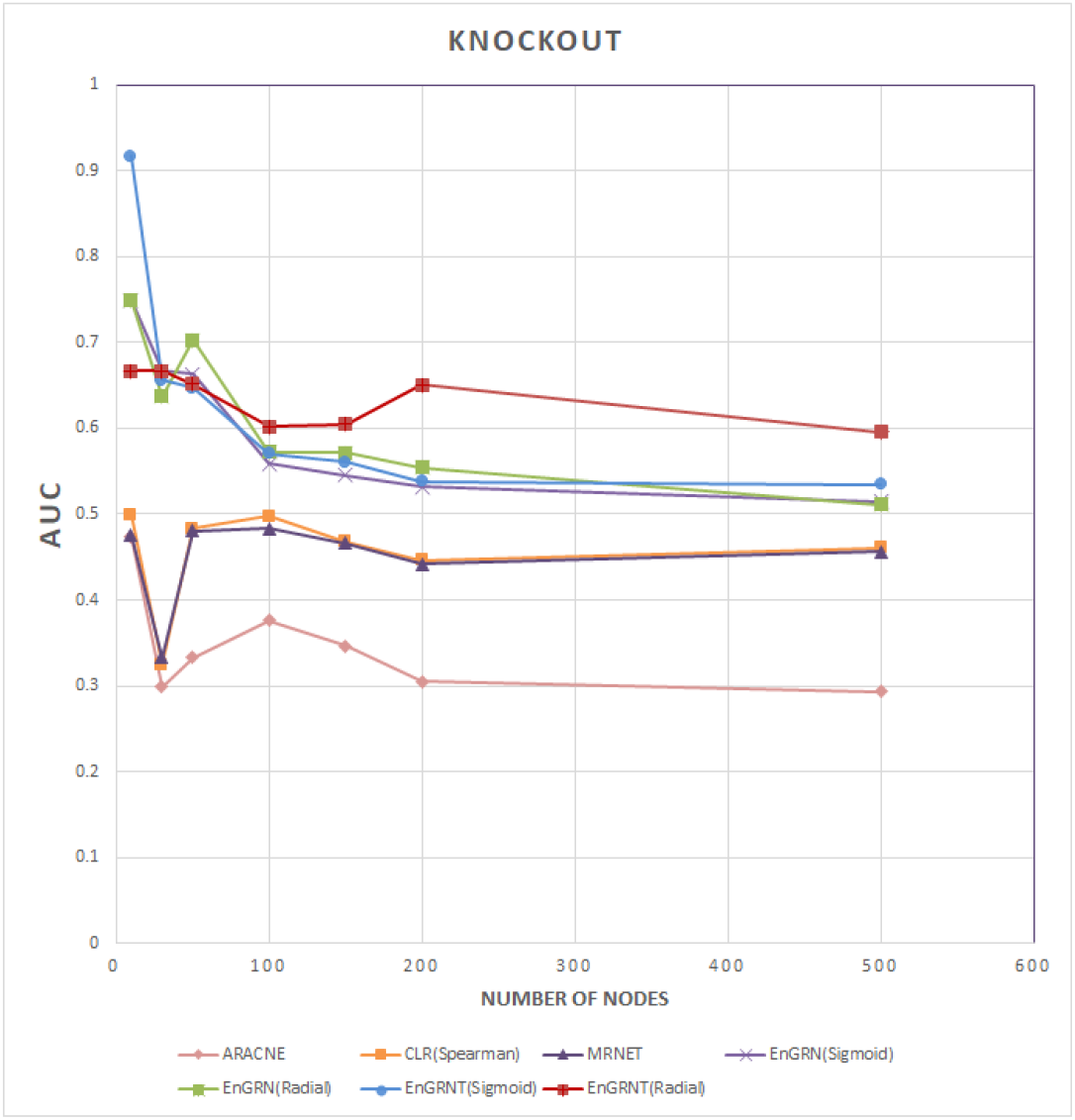
Prediction accuracy (AUC) of unsupervised and supervised methods on knockout experimental condition

**Fig.5.**
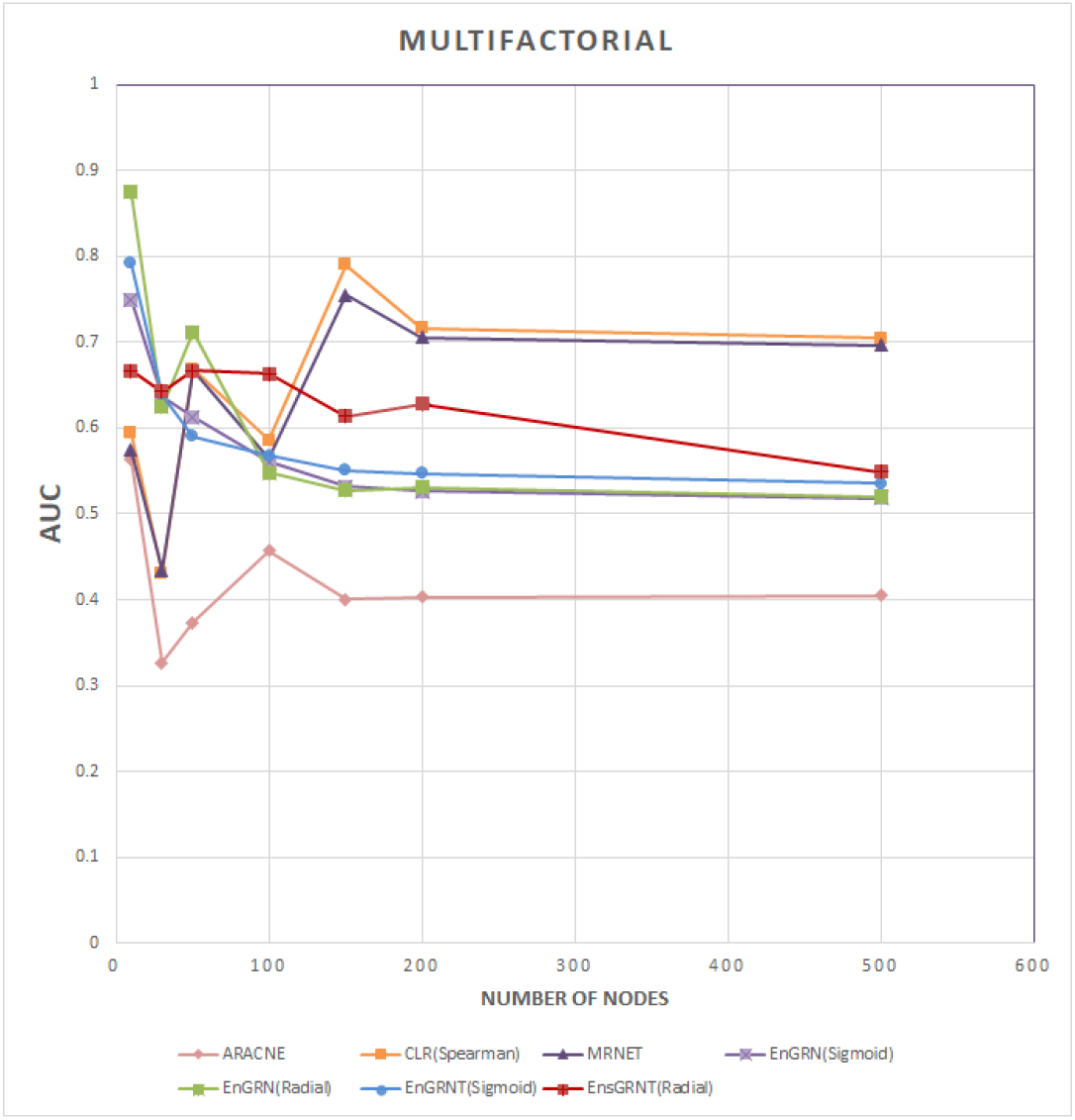
Prediction accuracy (AUC) of unsupervised and supervised methods on multifactorial experimental condition

These results indicate that employing the ensemble methods to train SVMs with proposed topological features in combination with gene expression data is well performed. Except for the multifactorial experimental conditions, the proposed method outperforms the unsupervised methods. In the case of small nodes (<30), the extraction of topological features was not very effective to increase the performance of the inference method and this is due to simple topology and the lack of nodes with the desired topological values, but in larger networks, the application of topological features improved prediction performance.

## 5. Conclusion

Reconstructing the structure of gene regulatory networks (GRNs) is an important task in the field of bioinformatics which could have profound impacts on understanding vital processes occurring in living organisms. The current machine learning-based methods can be divided in three groups which are supervised learning, unsupervised learning, and semi-supervised learning approaches. EnGRNT is categorized in supervised learning methods which transforms GRN inference problem to binary classification problem for each transcription factor. Based on this viewpoint, we are dealing with imbalanced classification problem. This is due to the fact that the number of regulatory relationships known between TF and target genes (Minority class) is much less than the number of regulatory relationships determined between TF and the target gene does not exist (Majority class). This despite the fact that current methods have ignored this concern which leads to be biased to the majority class with low accuracy for the minority class [11]. In order to tackle this problem, we exploited the Underbagging ensemble method that provides several bootstraps for each TF. Moreover, EnGRNT employs topological features to improve the accuracy of link prediction in the reconstructed GRNs.

The results of this work confirmed previously reported reports that in most cases, supervised learning methods have higher prediction accuracy than other methods, but when the number of nodes exceeds 150, unsupervised methods (CLR and MRNET) performed better than supervised methods in multifactorial experimental conditions. It was also observed that a large number of iterations in networks of different sizes were needed to estimate the prediction accuracy of a method.

The most important observation of this assessment is that there is no general way to infer GRNs in all biological conditions. On average, unsupervised methods with the exception of the multifactorial experimental conditions can achieve low accuracy. Our experimental results showed that the application of topological features can be effective in increasing the prediction accuracy of GRN inference methods. In summary, EnGRNT can be used to infer GRNs with acceptable accuracy for networks (<150) nodes using Gaussian kernel in experimental conditions (knockout, knockdown, and multifactorial). For large networks, it is critical to consider biological conditions for selecting an appropriate algorithm. We have adopted default parameter settings for the machine learning algorithms studied in the paper. We also plan to explore other parameter settings (e.g., different kernels with different parameter values in SVMs) in the future. In the present work, we proposed a supervised learning-based approach named as EnGRNT to GRN inference. The proposed method consists of two phases. First, the GRN inference is considered as a binary classification problem which performed for each TF individually. Second, a set of topological features are used to increase the accuracy of GRN inference. By evaluating the results of EnGRNT on simulated datasets, we found that the proposed approach could effectively overcome the challenge of imbalance data, and could provide promising results compared with conventional methods. As demonstrated in a series of recent publications making the methods freely available is the main trend in developing practically useful models or methods, the source code of EnGRNT is freely available at https://github.com/Khojasteh-hb/EnGRNT

## Acknowledgement

We would like to acknowledge the help that received from our colleagues in Machine Learning and Bioinformatics Laboratory (MLBL) of University of Zanjan, Zanjan, Iran.

